# Institutional Differences in the Stewardship and Research Output of United States Herbaria

**DOI:** 10.1101/2021.01.07.425759

**Authors:** Alexis Garretson

**Affiliations:** School of Systems Biology, George Mason University; Department of Biology, George Mason University

**Keywords:** data stewardship, herbaria, digitization, museums, natural history collections

## Abstract

Public policy decisions regarding institutional frameworks that govern the stewardship of biodiversity data at public and private institutions are an area of increasing importance. Museums, government agencies, and academic institutions across the United States maintain collections of biological specimens and information critical to scientific discovery. One subset of these natural history collections are herbaria, or collections of preserved plant matter and their associated data. In this study, I evaluate the current state of the digitization and databasing of herbariums contributing data to the SEInet Regional Network of North American Herbaria, and assess the impact of characteristics, particularly institution type (cultural sector institutions, public universities, private universities, or public land institutions), on the metrics of herbaria richness, digitization, and research usage. The results of this study suggest that institution type is significantly associated with the size, diversity, and digitization efforts of a herbarium collection. Specifically, cultural sector institutions tend to have larger and more diverse collections, followed by public and private universities, and finally public land institutions. Additionally, as herbarium size and richness increases, the research output of associated staff also increases. These results highlight that some institutions, particularly larger institutions located at universities or cultural sector institutions, may be better supported in the curation, stewardship, and digitization of large collections, allowing long-term access to the associated biodiversity data. Smaller institutions at public land institutions may need additional support in these endeavors, and may represent an area of unmet needs for digitization and curatorial funding and resources.

## Introduction

Biodiversity information is considered a complex good, having aspects of both common pool resources and public goods (Gómez-Zapata, Espinal-Monsalve, and Herrero-Prieto 2018; Dedeurwaerdere 2006; Escribano, Galicia, and Arino 2018). According to the conventional economic thought, both common pool resources and public goods are subject to overutilization and underprovision, theoretically leading to a market failure that requires government actors to support production and limit free riding (Anderson and Libecap 2014; Suarez and Tsutsui 2004; Escribano, Galicia, and Arino 2018; Hardin 1968; Samuelson 1954). Empirical research has demonstrated that under certain circumstances, private mechanisms can facilitate the provision of public goods, and can sometimes outperform government provision of public goods (Candela and Geloso 2018; Skarbek 2011, 2016; Stringham 2015). In addition, the work of Elinor Ostrom demonstrates that the tragedy of the commons can be averted and common pool resources can be efficiently managed for long-term use without requiring top-down government intervention (Poteete, Janssen, and Ostrom 2010; Ostrom 1990; Ostrom et al. 1994).

There is also tension within the economic literature between the efficiency of polycentric and centralized systems for the provision of public goods and services. Many scholars argue that polycentric institutions can better solve collective action problems because they are more flexible and can better incorporate local knowledge (Aligica and Tarko 2012; McGinnis 2000; Ostrom 2008; Coyne and Lemke 2011). Tarko (2015) suggests that a polycentric framework is part of what contributes to the success of the global scientific community. However, many of these authors recognize that polycentric institutions do not benefit from the economies of scale that can arise from more centralized approaches (Nagendra and Ostrom 2012; Thiel 2017; Ostrom 2010). This suggests that the provision of biodiversity data requires understanding the appropriate scale for the different natural history collection management activities, including specimen collection, long-term stewardship, and research utilization.

Herbaria represent a subset of natural history collections dedicated to the preservation of botanical specimens and dried plant matter (James et al. 2018). According to the Index Herbariorum 2019 report, there are 3,324 active herbaria containing 392,353,698 specimens (Thiers 2019). An herbarium specimen typically consists of exsiccatae - dried plant material mounted on paper with labels indicating key metadata, including date, species identification, and location. In general, herbarium specimens and their associated data are persistent, verifiable, and repeatable (James et al. 2018; L. M. Page et al. 2015; Holmes et al. 2016). Traditionally these specimens have served as critical sources of data to define a biological community at a point in time or for systematics and taxonomy studies to clarify and define species boundaries (L. M. Page et al. 2015; Besnard et al. 2018). However, expanding digitization efforts enables open access to biodiversity data and allows for big data investigations that would have previously been impossible (Devictor and Bensaude-Vincent 2016).

Presently, the best practices for digitization include not only the databasing of critical specimen metadata, but also the georeferencing and imaging of specimens (L. M. Page et al. 2015). Databasing is typically text-based metadata about an item, the date of collection, the date of taxonomic identification, the identities of the collector and the individual identifying the object, the taxonomic identification of the specimen, and locality information, typically including the county and state of collection, and occasionally including information about soil type, habitat, and biological community (Iwanycki 2009; Heidorn and Wei 2008). Often, the databasing step of digitization includes the assignment of a unique identifier, which may be a collection-specific barcode or a persistent identifier like an IGSN (Hobern, Hahn, and Robertson 2018; Nelson et al. 2015). Georeferencing involves using the text locality information to provide a mapped point location and associated error or uncertainty for the specimen to enable geoanalysis (Murphey et al. 2004). Imaging flat herbarium specimens typically includes the capture, processing, and archiving of a 2D image of the specimen (Nelson et al. 2015; Giraud et al. 2018). Imaging occasionally also includes 3D scans of structures such as fruit or buds of a specimen (Schneider et al. 2018), or microscopic images of pollen or other structures (Allan et al. 2019; Carranza-Rojas et al. 2017). Increasingly, digitization also includes genomics or phylogenetics data related to the specimen, and the digitization best practices may soon include standardized sequencing of specimen DNA (R. D. M. Page 2013; Taylor and Swann 1994; Leavitt et al. 2019).

The availability of digitized specimen data enables biodiversity big data research that was historically impossible. For example, georeferenced specimens can serve as baseline data for defining the impacts of climate change, invasive species, and other anthropogenic changes on plant communities (Lang et al. 2019; Dyderski et al. 2018; Ahern et al. 2010). Imaged specimens allow for studies of plant growth forms and coloration without requiring travel or loan of specimens and also provides a labeled training dataset for deep learning and taxonomic classification (Jimenez-Mejias, Cohen, and Naczi 2017; Collins et al. 2018). These images also contain information about the plant life stage at the time of collection, allowing researchers to assess changes in plant phenology in response to climate change and land use change (Cleland et al. 2007; Everill et al. 2014; MacGillivray, Hudson, and Lowe 2010). Additionally, advances in genomics technology can allow researchers a window into genetic changes and how species are distributed across a landscape through the sequencing and genetic profiling of collection specimens (Cozzolino et al. 2007; Konrade, Shaw, and Beck 2019; Snyman et al. 2018). Digitizing collections and making data publicly available should allow researchers to expand research usage of these collections, but these outputs may be dependent on particular management and institutional approaches to enabling herbarium research.

Much of the digitization progress in the United States has been funded through federal grants, particularly the Advancing Digitization of Biological Collections (ADBC) program, established by the National Science Foundation in 2011. ADBC provides funding to organizations to improve access to digitized specimens in US natural history collections (National Science Foundation 2015). This program established iDigBio, a centralized organization that coordinates the integration of the digital data resulting from digitization projects (Paul et al. 2013; Matsunaga et al. 2013; Nelson 2014). ADBC also supports Thematic Collections Network (TCN), which provides funding to networks of institutions with a shared strategy to digitize specimens from a specific research theme, such as a taxonomic focus or geographic region (Nelson 2014; National Science Foundation 2015). Eligible institutions include two- and four-year public and private colleges and universities; non-profit, non-academic cultural institutions including museums, botanical gardens, research labs, and professional societies; and state and local governments. There are presently more than 124,858,708 specimen records, 39,689,496 media records, and 1,623 record sheets aggregated on the iDigBio portal resulting from funding to 925 collections at 317 institutions.

<Table 1 near here> Different types of natural history institutions have different challenges in the process of managing and digitizing their collections (Mayernik et al. 2020). Smaller herbaria often struggle with prohibitively small budgets and few curatorial staff to assist in collection management and digitization tasks (Snow 2005; Harris and Marsico 2017). Herbaria located at larger universities may have a greater number of affiliated researchers, but herbarium management is rarely their primary role, and their particular research area may not meaningfully contribute to specimen collection or curatorial tasks (Feeley and Silman 2011).

**Table 1.**
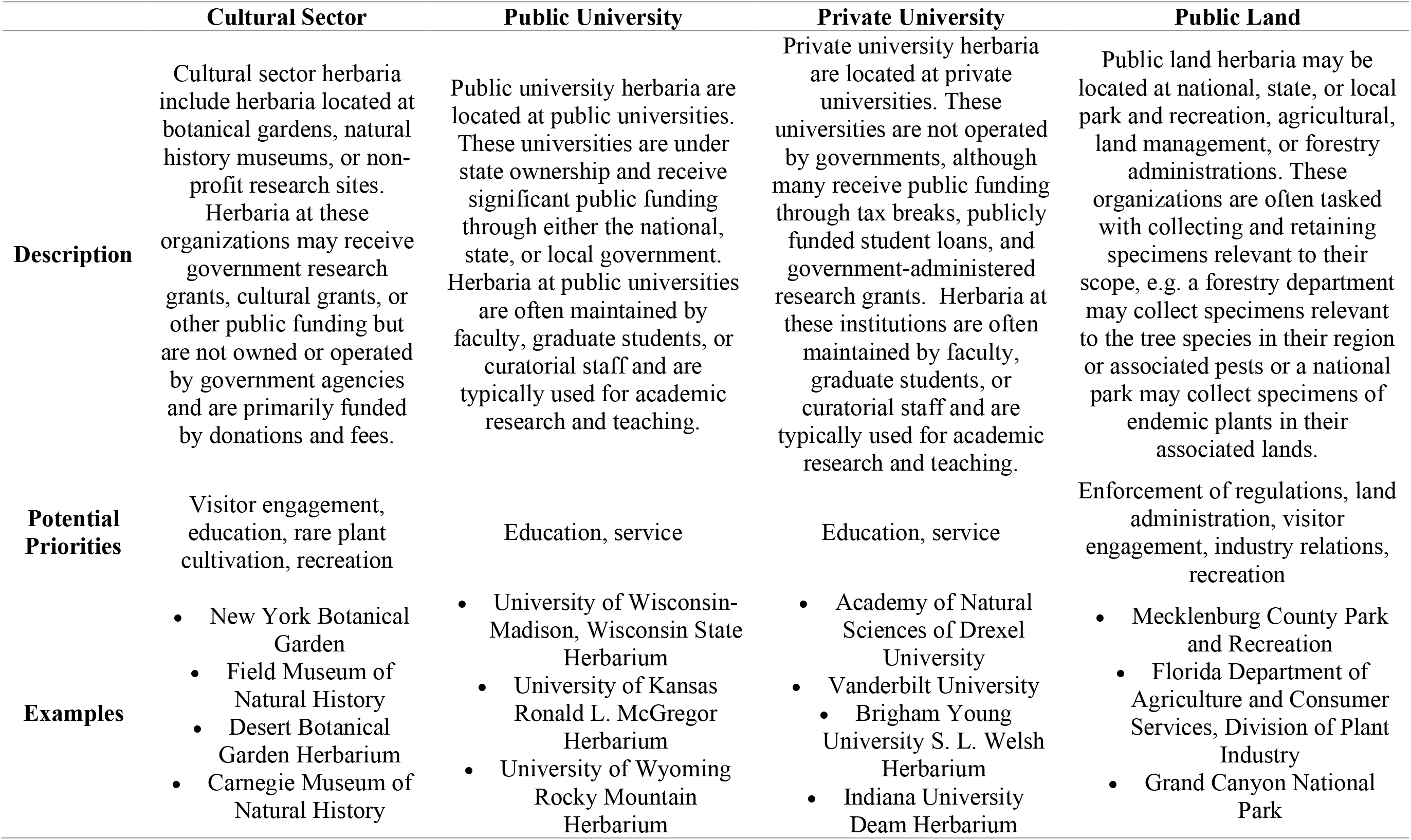
Table summarizing the features, potential institutional priorities, and examples of the institutional types discussed.

Cultural sector herbaria are often co-located with botanical gardens, field research sites, or natural history museums. The shared goals of cultural institutions include serving the public good, attaining financial stability, and supporting staff (Selwood 1999; Giardina and Rizzo 1994; Falk and Dierking 2008). Serving the public good takes a variety of forms, including serving as storehouses of cultural and scientific information, supporting research work on collection holdings, supporting social impact, and providing educational opportunities to the public (Scott 2006; Stanziola 2008). These institutions can vary in size, but their herbarium staff members are more likely to have curatorial tasks and taxonomic assessment as their primary role compared to university staff members. Variations in the curatorial role types and number of curators may have significant impacts on not only the size and richness of a herbarium, but also the progress made on digitization and incorporation into public databases. This ultimately can affect the research output associated with an individual herbarium.

Herbaria located at public land institutions face unique challenges compared to those at academic institutions. Namely, The Sundry Civil Act of March 3, 1879 (20 U.S.C. 59), requires that all physical object collections, including herbarium specimens, must eventually be archived in the Smithsonian Museum of Natural History. This means that, although a significant proportion of the collections are held by other organizations, like the Bureau of Land Management, the United States Geological Survey (USGS), the National Parks Service, and the Fish Wildlife Service, federally-managed collections are not typically locally managed or archived for long-term stewardship. The lack of accountability for physical objects has led to significant criticism from the scientific and data management community, particularly of the USGS (Office of the Inspector General 2017; Ruch 2018, 2019). A 2018 report highlighted that the agency lacked a policy for biological specimens, and its geological specimen policy had confusing language that could leave specimens at risk of destruction (Ruch 2018). In September of 2019, the USGS released their Policy on Scientific Working Collections (United States Geological Survey 2019), and though these policy changes are significant improvements, there are still concerns that these changes will not address risks to natural history collections under their administration (Ruch 2019).

Natural history collections at public lands are rarely collected exclusively for research purposes, and digitization for public research may not be a priority at these institutions. Additionally, the requirement that all physical objects collected by government entities must be eventually deposited into the Smithsonian Institution National Museum of Natural History for long-term archiving may leave publicly managed natural history collections without a strong incentive to manage their specimens for long-term research usage, as the specimens are not yet housed in their long-term home. In theory, requiring public land institutions to deposit specimens into the Smithsonian should enable research access and improve stewardship of federally collected physical objects. In practice, however, this policy leads to neglect and insufficient stewardship of the distributed working collections at the USGS and other institutions and often leaves specimens at risk of destruction or degradation due to storage under suboptimal conditions (Ruch 2018).

These academic, government-run, and cultural sector institutions across the world maintain substantial natural history collections containing biological specimens and physical objects (Suarez and Tsutsui 200). Because these institutions vary in their size, taxonomic diversity, and research goals, there are significant differences in management priorities. The variety of institutional frameworks under which natural history collections are managed provide both challenges and opportunities to researchers who rely on this data to produce scientific knowledge. The purpose of this study is to assess how the institutional management type impacts the size and richness of an herbarium, the progress towards digitization, and the research output.

## Methods

### Data Collection

Herbaria across North America submit data to the SEINet Regional Networks Of North, I extracted collection-level statistics for a total of 399 submitting institutions on February 21, 2020 using the SouthEast Regional Network of Expertise and Collections portal. This provided aggregated measures of herbarium size: the total number of specimens and the total number of specimens identified to species level. This also included measures of digitization efforts in the collection including the total number and percentage of specimens that are georeferenced or imaged. Additionally, the portal reports measures of herbarium taxonomic diversity including: number of families, genera, species, and total taxonomic groups including in the collection. Finally, the portal also reports the total number of type specimens, or specimens that have permanent taxonomic designations used by other researchers to confirm identifications and define taxonomic boundaries, in the collection. Herbaria outside of the United States were excluded from this analysis, and each remaining herbarium was classified as either associated with a public land institution, public university, private university, cultural sector institution, or for-profit organization.

In addition to measures of herbarium richness, five measures of research activity were collected for each herbaria. First, the total number of research staff for each institution was collected through the Index Herbariorum from the New York Botanical Gardens, which provides a list of associated staff members for each indexed herbarium. If an institution did not have an associated staff list, the homepage of the herbarium was used as a source of the number of staff members. For each collection, the total number of Google Scholar search results from a search for the institution name was recorded as a proxy for the number of times the institution name appears in publications, either as an author affiliation, in the acknowledgments, in research methods, or in cited material. This is admittedly a coarse measure of research output, so additionally, each staff member affiliated with the herbarium institutions was screened for a Google Scholar page. The total number of research articles and associated citations for each of the staff members with a Google Scholar page was recorded.

### Statistical Analysis

To determine the correlations between the variables of interest, the Pearson correlation was calculated for each of the pairwise comparisons. Additionally, a one-way ANOVA was used to determine the association between the institution type and measures of herbarium research output and richness. In outcome variables with an ANOVA significant at *P* < 0.05, a Tukey’s HSD test was used to determine the significant differences between the mean values of institutions and statistically significant groupings between the institution types. Finally, the herbarium richness measures were compared to research outcome variables using linear regressions. All data analysis was performed in R (version 3.6.2) with RStudio (version 1.1.453).

## Results

### Summary of data collection

<Table 2 near here> Of the 339 total institutions that submit data to SEInet, 28 were omitted from the present study for having fewer than 10 databased specimens. A further 10 were omitted for being outside of the United States, leaving 301 institutions included for further analysis. After categorizing by type, the majority of the herbaria are housed at universities, with 52 at private universities and 174 at public universities. There are a total of 30 institutions that were categorized as cultural sector institutions, primarily housed in botanical gardens and museums. A total of 36 institutions are categorized as public land institutions, primarily housed at national parks, other federal institutions like the Bureau of Land Management and Forest Service, and state or local park departments. Finally, the remaining five herbaria collections are housed at fully private institutions, with two at for-profit companies and three at for-profit nature preserves. Due to the low number of representative collections, the fully private herbaria were excluded from future analysis by type, though the staff associated with the herbaria were included in analyses of research output and herbaria measures.

**Table 2.** Table of the distribution of herbarium richness variables and research productivity variables with the mean ± the standard deviation. A total of 292 institutions are included in the table, distributed across the institution types. If the ANOVA was significant, the superscripts in boxes designate the grouping results of a Tukey’s HSD test.

|  | **Cultural Sector**<br>**(n = 30)** | **Public University**<br>**(n = 174)** | **Private University**<br>**(n = 52)** | **Public Land**<br>**(n = 36)** | **ANOVA**<br>**p-value** |
| --- | --- | --- | --- | --- | --- |
| **Specimens** | 116,791.56 ± 223,471.76<sup>a</sup> | 54,479.36 ± 100,151.03<sup>b</sup> | 37,992.08 ± 79,263.06<sup>bc</sup> | 4,040.31 ± 7,125.46<sup>c</sup> | 0.0005*** |
| **Specimens Identified to Species** | 99,005.20 ± 202,174.01<sup>a</sup> | 45,742.58 ± 90,030.05<sup>b</sup> | 32,582.29 ± 72,599.88<sup>b</sup> | 2,975.19 ± 3,971.88<sup>b</sup> | 0.001** |
| **Families** | 217.37 ± 104.98<sup>a</sup> | 169.72 ± 110.43<sup>a</sup> | 162.92 ± 85.62<sup>a</sup> | 97.64 ± 63.39<sup>b</sup> | <0.0001*** |
| **Genera** | 1,569.87 ± 1,529.01<sup>a</sup> | 975.21 ± 912.87<sup>b</sup> | 830.60 ± 712.88<sup>bc</sup> | 404.39 ± 380.52<sup>c</sup> | <0.0001*** |
| **Species** | 7,402.43 ± 10,628.58<sup>a</sup> | 3,488.98 ± 4,042.96<sup>b</sup> | 2,787.13 ± 3,515.27<sup>bc</sup> | 908.17 ± 1,099.48<sup>c</sup> | <0.0001*** |
| **Total Taxa** | 8,609.27 ± 12,476.82<sup>a</sup> | 4,015.69 ± 4,768.41<sup>b</sup> | 3,150.06 ± 4,256.54<sup>bc</sup> | 1,076.24 ± 1,183.02<sup>c</sup> | <0.0001*** |
| **Types** | 17,627.90 ± 83,965.76<sup>a</sup> | 121.96 ± 630.07<sup>b</sup> | 175.10 ± 809.22<sup>b</sup> | 0.68 ± 2.61<sup>b</sup> | 0.011* |
| **Specimens Georeferenced** | 40,810.50 ± 64,715.56<sup>a</sup> | 18,034.30 ± 61,907.48<sup>ab</sup> | 7,661.46 ± 23,378.49<sup>b</sup> | 795.42 ± 1,347.14<sup>b</sup> | 0.013* |
| **Percent of Specimens Georeferenced** | 48% ± 36%<sup>a</sup> | 26% ± 33%<sup>ab</sup> | 28% ± 36%<sup>ab</sup> | 36% ± 44%<sup>b</sup> | 0.01** |
| **Specimens Imaged** | 51,594.00 ± 156,336.23 | 33,382.05 ± 65,320.47 | 33,104.92 ± 76,464.21 | 2,712.44 ± 6,941.53 | 0.73 |
| **Percent of Specimens Imaged** | 48% ± 46% | 58% ± 44% | 66% ± 43% | 45% ± 49% | 0.13 |

Across the institutions, a total of 1,102 staff members were found across all herbaria, because some herbaria shared staff members, this led to 1,024 unique staff members. Of these staff members, 24.31% (268) had active Google Scholar profile pages (Table 3). The distribution of herbaria richness across the remaining types (cultural sector institution, public university, private university, and public land institution) is presented in Table 1.

**Table 3.**
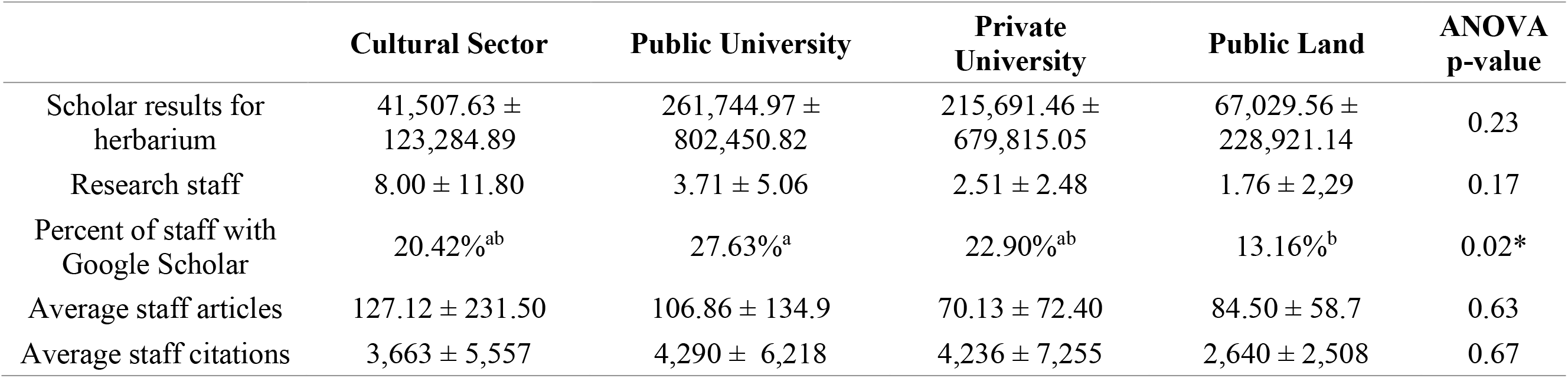
Results of the research assessments for each herbaria. Reported average number of citations and average number of articles are for researchers with a Google Scholar page.

### Cultural Sector Institutions Have Larger And More Diverse Collections

<Figure 1 near here> Measures of herbarium richness are positively correlated with each other, including measures of size, diversity, and digitization (Figure 1, Table 3). However, the percent of imaged specimens is negatively correlated with both the percent of georeferenced specimens (r = −0.21, *P* = 0.0001) and the total number of georeferenced specimens (r = −0.11, *P* = 0.04). Additionally, the number of research staff at a herbarium is significantly and positively correlated with measures of herbarium richness, except for the number of type specimens in the institution (Figure 1). In every herbarium richness, diversity, and digitization analysis, public lands were in the Tukey grouping with the lowest mean, indicating public lands tended to have smaller and less diverse collections (Table 2). In contrast, cultural sector institutions were in the highest group, indicating larger and more diverse collections. In most assessments, the universities, both public and private, are in an intermediate group. However, in comparing within universities, public universities tended to have higher mean average metrics of diversity than private universities (Table 2).

**Figure 1.**
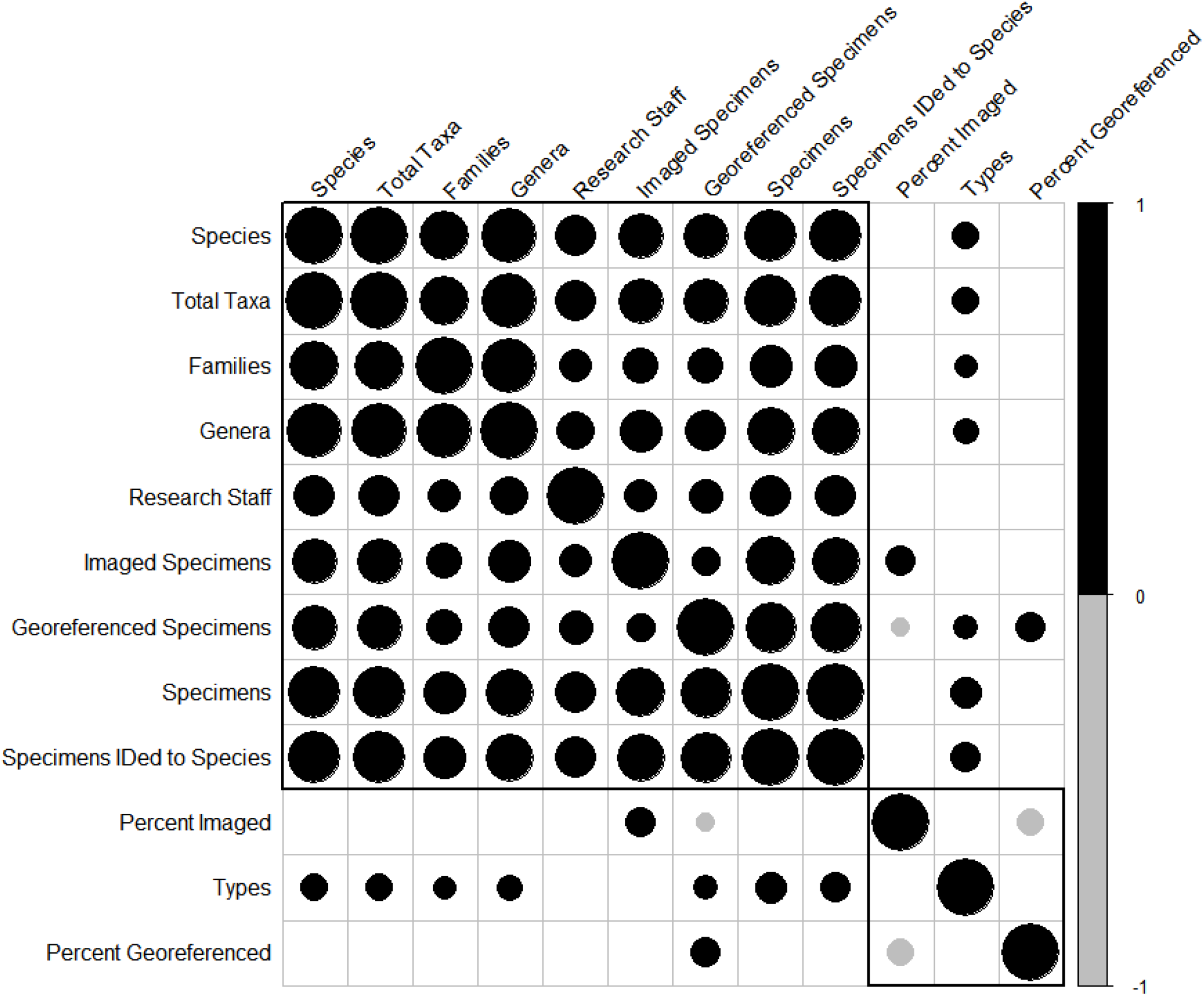
Pearson’s correlation between measures of herbaria richness, with more positive values indicated by darker blue and more negative values indicated by darker red. Asterisks indicate correlations significant at *P* < 0.05.

### Staff At Larger And More Diverse Herbaria Publish More Research Articles

<Figure 2 near here> No significant differences were found between institution type and measures of collection research output. However, there is a significant and positive relationship between the number of articles published by staff with Google Scholar pages and measures of herbaria richness (Figure 2, Table 3). Particularly, the number of articles by staff members increased with the total number of the research staff at their home institution (b = 4.11, *P* < 0.0001, Figure 2A). This relationship also holds between the total Google Scholar results for the researcher name in the absence of a Google Scholar page (b = 3.77, *P* < 0.0001). There is also a significant positive correlation between a number of herbaria richness measures and the number of research articles of research staff with Google Scholar pages, namely the number of collection specimens (*P* = 0.0054), the total taxonomic groups represented by the collection (*P* = 0.03), the number of species groups in the collection (*P* = 0.04), and the number of type specimens (*P* = 0.03) (Figure 2). Finally, there was a significant difference in the percentage of staff associated with each institution type that had a Google Scholar profile page (Table 3).

**Figure 2.**
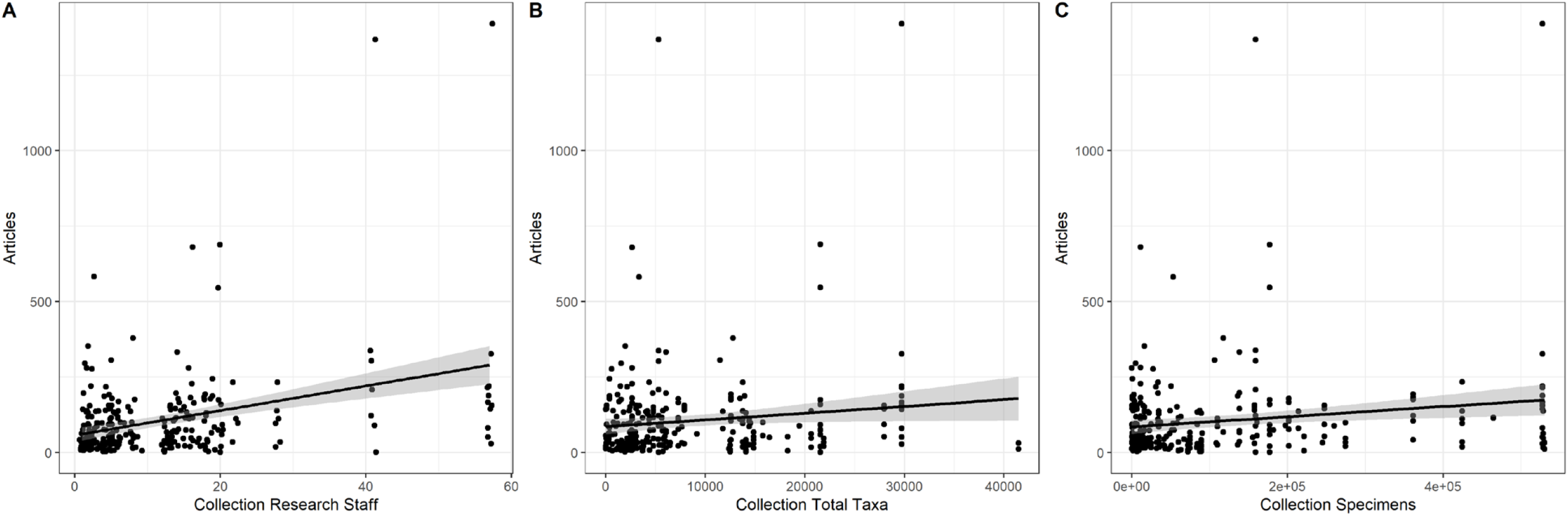
Plots of the number of articles published by herbaria staff with Google Scholar pages plotted against the (A) total number of research staff at the collection they work at, (B) the total taxa represented by collection specimens, and (C) the total number of specimens in the collection. Linear regression plotted with standard error and jitter added to points to prevent overplotting.

## Discussion

### Larger Collections Tend To Be More Diverse

One of the findings of this analysis is that herbarium richness in one area is positively correlated with herbarium richness in other measures. As size increases, the diversity, and digitization of the collection also increases. The institution type is significantly related to the institution type, with richer and larger herbaria at cultural sector institutions, followed by private universities, public universities, and finally public land institutions. Consistently, public land institutions lag behind both cultural sector institutions and university institutions across richness measures, suggesting management features of public land collection institutions may not lend themselves to large and diverse collections.

Cultural sector institutions outperform other institution types in the size and diversity of their collections. This may be because a greater overall research activities occurring at these institutions, as cultural sector institutions were found to have a higher number of type specimens --an indicator of active systematics research-- and higher percent of specimens georeferenced -- an indicator of active big data diversity research and ongoing digitization-- in their collections (Table 1, Table 2). Alternatively, for this relationship may be because cultural sector institutions are often co-located with botanical gardens (living collections), leading to easy access to exotic plants that may otherwise be difficult to collect as specimens. While this would increase the number of unique species in the collection, thus increasing the associated biodiversity, this may not adequately support the needs of researchers interested in documenting wild species occurrence and differences across a species’ natural range. Increasingly, cultural institutions like museums and botanical gardens also harness visitor enthusiasm for digitization tasks through citizen science tools, which can support digitization activities (Garretson et al. 2020; Garretson Forthcoming). Additionally, staff at cultural sector institutions are often not subject to other non-curation tasks, while collectors at public lands may have obligations to collect specific specimens relevant to survey work, and collectors at universities may have additional restrictions on time, including teaching, mentorship, and service (Snow 2005; Ab Rahim et al. 2013; Adams and Griliches 2000).

However, the negative correlation between the percent of specimens that are imaged and the percent of specimens georeferenced may suggest that there is a tradeoff between digitization task types (Figure 1). There are very different skill sets and technological requirements to georeference specimens compared to imaging specimens (Nelson et al. 2015; Blagoderov et al. 2012). Standardized imaging of specimens requires at minimum, a high-quality digital camera, lighting fixtures, a camera stand, a color scale, and a ruler. This setup can cost upwards of US$ 1500, which may be prohibitive for a small herbarium (Harris and Marsico 2017). Therefore, institutions with a limited budget may need to focus on only one digitization task at a time, and may focus on georeferencing due to the financial investment required to appropriately image specimens. As digitization tasks expand to include generating genetic data for each specimen, the financial and technological needs to meet best practices of digitization will grow, increasing the disparity between institutions in digitization completion.

### Research output and herbaria richness

The lack of association between institution type and individual researcher outcomes may be because there are too many confounding factors like career stage, position type, or age (Beaudry and Allaoui 2012; Gonzalez-Brambila and Veloso 2007; Wang et al. 2017). There may also be differences in the subset of researchers across all institution types that are likely to have active Google Scholar accounts. Particularly, younger researchers may be more likely to have an active online presence, as previous studies have found the age and career stage of a researcher can impact their assessment of the importance online academic presence (Mierzecka, Kisilowska, and Suminas 2020; Arshad and Ameen 2017; Wang et al. 2017). This may mean that the dataset of researchers used in this study may not be a random sample of researchers at the institutions. However, a positive relationship between herbaria measures and research outcomes means that a larger dataset could reveal significant institutional trends. The association with the number of staff might mean that collaborations and staff community could lead to more research, suggesting that larger institutions may generally have greater research output for any given faculty. This is in line with prior studies of research output of university faculty members that have suggested that larger public universities have greater research output compared to that of smaller private universities (Ab Rahim et al. 2013; Gonzalez-Brambila and Veloso 2007).

### Conclusions

These results demonstrate that there are institutional differences in collection stewardship and there is a link between institutional type and its effectiveness in contributing to publicly available biodiversity data. The results of this study show that institution type is significantly associated with the size, diversity, and digitization of a herbarium collection. However, past studies have found that critical occurrence records and rare taxa can be found in small natural history collections (Glon et al. 2017). Small and distributed herbaria may be better able to catalogue and collect unique local flora, but may not have the resources to support larger research endeavors. There are also certainly differences in the institutional priorities across all institutions, particularly in the mission, vision, and objectives of the institutions which can drive data collection methods, focuses, and digitization protocols. Additionally, the results demonstrate that as herbaria richness increases, the research output of associated staff increases in kind. This suggests that there may be economies of scale in research output because as the number of researchers grow, research output per researcher grows accordingly. Decentralized, small collections can benefit from local knowledge and unique records, but centralization has significant returns to scale in collection stewardship, digitization, research output.

### Implications of differences between cultural sector and public land institutions

While small herbaria can be important to researchers due to their ability to add unique occurrence information for rare plants (Glon et al. 2017), these findings suggest that smaller herbaria, particularly those located at public land institutions, may be under digitized and under incorporated into the publicly available biodiversity datasets. This may be because smaller herbaria tend to have fewer staff and fewer resources to support data collection, digitization, and access (Harris and Marsico 2017; Snow 2005; Blagoderov et al. 2012). Larger herbaria benefit from scale with respect to personnel, access to technology, and curatorial skill sets that may lead to greater opportunities for collaborative publication, collection research usage, and expansion of the collection. This demonstrates that smaller herbaria can act as polycentric institutions - better able to incorporate local knowledge and particularities of their surrounding region while other aspects of collection curation, namely digitization and long term storage, is better suited to more centralized archives. This mismatch between the institutions and researchers best positioned to collect novel specimens and institutions best positioned to digitize and manage the data in the long term represents an ongoing challenge in the management of small collections. This may also represent an important area for increased investment in supporting smaller institutions in building the capacity to support additional digitization and collection activities.

This finding supports the policy requiring federal entities to plan for submission of objects to the Smithsonian Institution. The Smithsonian Museum of Natural History is a public-nonprofit partnership, and contains more than 156 natural history specimens. The Smithsonian Museums benefit from economies of scale and often are the first institutions to have access to cutting edge curatorial technology including 3D scanners and genomics tools. However, the challenge in implementing this policy, particularly in the USGS, has been that this policy leaves USGS staff without significant investment in the long-term stewardship of the physical objects in question, and often without the training to appropriately store the items to prevent deterioration and destruction. Requiring the development of collection management plans may help address some of these concerns, but may not be sufficient to ensure proper management of the collection items before they reach the Smithsonian Institution. Like the USGS, many small herbaria need to maintain transition plans for their collections in the event that they lose funding, complete the associated research project, or face other risks to the datasets (Mayernik et al. 2020). Tools like persistent identifiers, such as IGSNs, can help prevent some of these hurdles by ensuring critical metadata is associated with natural history specimens throughout its lifecycle (Hobern, Hahn, and Robertson 2018).

### Directions for Future Research

Herbaria are only one type of biodiversity information facility, so assessing whether these trends hold with museums, field stations, and published biodiversity data and literature may be key to further understanding how research institution types and digitization efforts impact biological discovery and knowledge generation. Other studies have suggested that substantial differences exist between universities with and without a medical school, and that these institutional differences may be more important than the private/public differentiation (Ahn, Charnes, and Cooper 1988). Finer-grain differences in the institutional type might also be relevant for the understanding of how institutional type might influence the collection richness and research effort, so future studies should consider the age of the institution and its operating budget.

Better understanding the institutional differences in the digitization of natural history collections can assist in targeting collections that might need additional support through granting agencies, like the ADBC and can improve our understanding of how policies regulating the deposition of federally collected specimens might impact long term collection outcomes. Collections of natural history specimens are an invaluable resource in understanding scientific processes, and contribute to critical research in policy-relevant areas like public health and pandemic forecasting, monitoring of climate and ecological change, and basic biological research. Ensuring the ongoing protection, digital preservation, and research access to these items is a critical aspect of stewarding our research resources and better understanding our changing environment.

## References

Ab Rahim, Ina Suryani, Aizan Yaacob, Noor Hashima Abd Aziz, Salleh Abd Rashid, and Hazry Desa. 2013. “Research Publication Output by Academicians in Public and Private Universities in Malaysia.” International Journal of Higher Education 2 (1). Sciedu Press: 84–90.

Adams, James D., and Zvi Griliches. 2000. “Research Productivity in a System of Universities.” In The Economics and Econometrics of Innovation, edited by David Encaoua, Bronwyn H. Hall, François Laisney, and Jacques Mairesse, 105–40. Boston, MA: Springer US. doi:10.1007/978-1-4757-3194-1_5.

Ahern, Robert G., Douglas A. Landis, Anton A. Reznicek, and Douglas W. Schemske. 2010. “Spread of Exotic Plants in the Landscape: The Role of Time, Growth Habit, and History of Invasiveness.” Biological Invasions 12 (9): 3157–69. doi:10.1007/s10530-010-9707-x.

Ahn, Taesik, Abraham Charnes, and William W. Cooper. 1988. “Some Statistical and DEA Evaluations of Relative Efficiencies of Public and Private Institutions of Higher Learning.” Socio-Economic Planning Sciences 22 (6): 259–69. doi:10.1016/0038-0121(88)90008-0.

Aligica, Paul D., and Vlad Tarko. 2012. “Polycentricity: From Polanyi to Ostrom, and Beyond.” Governance 25 (2): 237–62. doi:10.1111/j.1468-0491.2011.01550.x.

Allan, E Louise, Laurence Livermore, Benjamin W Price, Olha Shchedrina, and Vincent S Smith. 2019. “A Novel Automated Mass Digitisation Workflow for Natural History Microscope Slides.” Biodiversity Data Journal 7 (March). doi:10.3897/BDJ.7.e32342.

Anderson, Terry L., and Gary D. Libecap. 2014. Environmental Markets: A Property Rights Approach. Cambridge Studies in Economics, Cognition, and Society. New York: Cambridge University Press.

Arshad, Alia, and Kanwal Ameen. 2017. “Scholarly Communication in the Age of Google: Exploring Academics’ Use Patterns of e-Journals at the University of the Punjab.” The Electronic Library 35 (1): 167–84. doi:10.1108/EL-09-2015-0171.

Beaudry, Catherine, and Sedki Allaoui. 2012. “Impact of Public and Private Research Funding on Scientific Production: The Case of Nanotechnology.” Research Policy 41 (9): 1589– 1606. doi:10.1016/j.respol.2012.03.022.

Besnard, Guillaume, Myriam Gaudeul, Sébastien Lavergne, Serge Muller, Germinal Rouhan, Alexander P. Sukhorukov, Alain Vanderpoorten, and Florian Jabbour. 2018. “Herbarium-Based Science in the Twenty-First Century.” Botany Letters 165 (3–4). Taylor & Francis: 323–27. doi:10.1080/23818107.2018.1482783.

Blagoderov, Vladimir, Ian J. Kitching, Laurence Livermore, Thomas J. Simonsen, and Vincent S. Smith. 2012. “No Specimen Left behind: Industrial Scale Digitization of Natural History Collections.” ZooKeys, no. 209: 133.

Candela, Rosolino A., and Vincent J. Geloso. 2018. “The Lightship in Economics.” Public Choice 176 (3): 479–506. doi:10.1007/s11127-018-0573-x.

Carranza-Rojas, Jose, Herve Goeau, Pierre Bonnet, Erick Mata-Montero, and Alexis Joly. 2017. “Going Deeper in the Automated Identification of Herbarium Specimens.” BMC Evolutionary Biology 17 (August). doi:10.1186/s12862-017-1014-z.

Cleland, Elsa E., Isabelle Chuine, Annette Menzel, Harold A. Mooney, and Mark D. Schwartz. 2007. “Shifting Plant Phenology in Response to Global Change.” Trends in Ecology & Evolution 22 (7): 357–65. doi:10.1016/j.tree.2007.04.003.

Collins, Matthew, Gaurav Yeole, Paul Frandsen, Rebecca Dikow, Sylvia Orli, and Renato Figueiredo. 2018. “A Pipeline for Deep Learning with Specimen Images in IDigBio -Applying and Generalizing an Examination of Mercury Use in Preparing Herbarium Specimens.” Biodiversity Information Science and Standards 2 (July): e25699. doi:10.3897/biss.2.25699.

Coyne, Christopher J, and Jayme S Lemke. 2011. “Polycentricity in Disaster Relief.” Studies in Emergent Order 4: 40–57.

Cozzolino, Salvatore, Donata Cafasso, Giuseppe Pellegrino, Aldo Musacchio, and Alex Widmer. 2007. “Genetic Variation in Time and Space: The Use of Herbarium Specimens to Reconstruct Patterns of Genetic Variation in the Endangered Orchid Anacamptis Palustris.” Conservation Genetics 8 (3): 629–39. doi:10.1007/s10592-006-9209-7.

Dedeurwaerdere, Tom. 2006. “The Institutional Economics of Sharing Biological Information.” International Social Science Journal 58 (188): 351-+. doi:10.1111/j.1468-2451.2006.00623.x.

Devictor, Vincent, and Bernadette Bensaude-Vincent. 2016. “From Ecological Records to Big Data: The Invention of Global Biodiversity.” History and Philosophy of the Life Sciences 38 (4): pUNSP 13. doi:10.1007/s40656-016-0113-2.

Dyderski, Marcin K., Sonia Paz, Lee E. Frelich, and Andrzej M. Jagodzinski. 2018. “How Much Does Climate Change Threaten European Forest Tree Species Distributions?” Global Change Biology 24 (3): 1150–63. doi:10.1111/gcb.13925.

Escribano, Nora, David Galicia, and Arturo H. Arino. 2018. “The Tragedy of the Biodiversity Data Commons: A Data Impediment Creeping Nigher?” Database-the Journal of Biological Databases and Curation, April, bay033. doi:10.1093/database/bay033.

Everill, Peter H., Richard B. Primack, Elizabeth R. Ellwood, and Eli K. Melaas. 2014. “Determining Past Leaf-out Times of New England’s Deciduous Forests from Herbarium Specimens.” American Journal of Botany 101 (8): 1293–1300. doi:10.3732/ajb.1400045.

Falk, John H., and Lynn D. Dierking. 2008. “Re-Envisioning Success in the Cultural Sector.”Cultural Trends 17 (4). Routledge: 233–46. doi:10.1080/09548960802615372.

Feeley, Kenneth J., and Miles R. Silman. 2011. “Keep Collecting: Accurate Species Distribution Modelling Requires More Collections than Previously Thought.” Diversity and Distributions 17 (6): 1132–40. doi:10.1111/j.1472-4642.2011.00813.x.

Garretson, Alexis. Forthcoming. “Citizen Science Can Improve Visitor Experiences and Research Outcomes in Museums and Cultural Institutions.” In What’s Emerging in the Field? Museum Computer Network.

Garretson, Alexis, Megan Napoli, Natalie Feldsine, Penelope Adler-Colvin, and Elizabeth Long.2020. “Vernal Pool Amphibian Breeding Ecology Monitoring from 1931 to Present: A Harmonised Historical and Ongoing Observational Ecology Dataset.” Biodiversity Data Journal 8 (April). Pensoft Publishers: e50121. doi:10.3897/BDJ.8.e50121.

Giardina, Emilio, and Ilde Rizzo. 1994. “Regulation in the Cultural Sector.” In Cultural Economics And Cultural Policies, edited by Alan Peacock and Ilde Rizzo, 125–42. Dordrecht: Springer Netherlands. doi:10.1007/978-94-011-1140-9_10.

Giraud, Michel, Quentin Groom, Ann Bogaerts, Sofie de Smedt, Hannu Saarenmaa, Noortje Wijkamp, Sarah Philips, and Zhengzhe Wu. 2018. “Best Practice Guidelines for Imaging of Herbarium Specimens.” Innovation and Consolidation for Large Scale Digitization of Natural Heritage, January, 41.

Gómez-Zapata Jonathan Daniel, Nora Elena Espinal-Monsalve, and Luis César Herrero-Prieto. 2018. “Economic Valuation of Museums as Public Club Goods: Why Build Loyalty inCultural Heritage Consumption?” Journal of Cultural Heritage 30 (March): 190–98. doi:10.1016/j.culher.2017.09.010.

Gonzalez-Brambila, Claudia, and Francisco M. Veloso. 2007. “The Determinants of Research Output and Impact: A Study of Mexican Researchers.” Research Policy 36 (7): 1035–51. doi:10.1016/j.respol.2007.03.005.

Hardin, Garrett. 1968. “The Tragedy of the Commons.” Science 162 (3859). American Association for the Advancement of Science: 1243–48. doi:10.1126/science.162.3859.1243.

Harris, Kari M., and Travis D. Marsico. 2017. “Digitizing Specimens in a Small Herbarium: A Viable Workflow for Collections Working with Limited Resources1.” Applications in Plant Sciences 5 (4). doi:10.3732/apps.1600125.

Heidorn, P. Bryan, and Qin Wei. 2008. “Automatic Metadata Extraction from Museum Specimen Labels.” In International Conference on Dublin Core and Metadata Applications, 57–68.

Hobern, Donald, Andrea Hahn, and Tim Robertson. 2018. “Options to Apply the IGSN Model to Biodiversity Data.” Biodiversity Information Science and Standards 2 (May). Pensoft Publishers: e27087. doi:10.3897/biss.2.27087.

Holmes, Michael W., Talisin T. Hammond, Guinevere O. U. Wogan, Rachel E. Walsh, Katie LaBarbera, Elizabeth A. Wommack, Felipe M. Martins, et al. 2016. “Natural History Collections as Windows on Evolutionary Processes.” Molecular Ecology 25 (4): 864–81. doi:10.1111/mec.13529.

Iwanycki, Natalie. 2009. “Guidelines for Collecting Herbarium Specimens.” Royal Botanical Gardens, July, 5.

James, Shelley A., Pamela S. Soltis, Lee Belbin, Arthur D. Chapman, Gil Nelson, Deborah L. Paul, and Matthew Collins. 2018. “Herbarium Data: Global Biodiversity and Societal Botanical Needs for Novel Research.” Applications in Plant Sciences 6 (2): e1024. doi:10.1002/aps3.1024.

Jimenez-Mejias, Pedro, James I. Cohen, and Robert F. C. Naczi. 2017. “The Study of Online Digitized Specimens Revalidates Andersonglossum Boreale as a Species Different from A-Virginianum (Boraginaceae).” Phytotaxa 295 (1): 22–34. doi:10.11646/phytotaxa.295.1.2.

Konrade, Lauren, Joey Shaw, and James Beck. 2019. “A Rangewide Herbarium-derived Dataset Indicates High Levels of Gene Flow in Black Cherry (Prunus Serotina).” Ecology and Evolution 9 (3): 975–85. doi:10.1002/ece3.4719.

Lang, Patricia L. M., Franziska M. Willems, J. F. Scheepens, Hernán A. Burbano, and Oliver Bossdorf. 2019. “Using Herbaria to Study Global Environmental Change.” New Phytologist 221 (1): 110–22. doi:10.1111/nph.15401.

Leavitt, Steven D., Rachel Keuler, Clayton C. Newberry, Roger Rosentreter, and Larry L. St Clair. 2019. “Shotgun Sequencing Decades-Old Lichen Specimens to Resolve Phylogenomic Placement of Type Material.” Plant and Fungal Systematics 64 (2). Sciendo: 237–47. doi:10.2478/pfs-2019-0020.

MacGillivray, Fran, Irene L. Hudson, and Andrew J. Lowe. 2010. “Herbarium Collections and Photographic Images: Alternative Data Sources for Phenological Research.” In Phenological Research: Methods for Environmental and Climate Change Analysis, edited by Irene L. Hudson and Marie R. Keatley, 425–61. Dordrecht: Springer Netherlands. doi:10.1007/978-90-481-3335-2_19.

Matsunaga, Andréa, Renato Figueiredo, Alex Thompson, Gregory Traub, Reed Beaman, and José AB Fortes. 2013. “Integrated Digitized Biocollections (IDigBio) Cyberinfrastructure Status and Futures.” In TDWG 2013 ANNUAL CONFERENCE.

Mayernik, Matthew S., Kelsey Breseman, Robert R. Downs, Ruth Duerr, Alexis Garretson, Chung-Yi (Sophie) Hou, and Environmental Data Governance Initiative (EDGI) and Earth Science Information Partners (ESIP) Data Stewardship Committee. 2020. “Risk Assessment for Scientific Data.” Data Science Journal 19 (1). Ubiquity Press: 10. doi:10.5334/dsj-2020-010.

McGinnis, Michael Dean. 2000. Polycentric Games and Institutions: Readings from the Workshop in Political Theory and Policy Analysis. University of Michigan Press.

Mierzecka, Anna, Małgorzata Kisilowska, and Andrius Suminas. 2020. “Researchers’ Expectations Regarding the Online Presence of Academic Libraries | Mierzecka | College & Research Libraries.” Accessed March 20. doi:https://doi.org/10.5860/crl.78.7.934.

Murphey, Paul C, Robert P Guralnick, David Neufeld, and J Allen Ryan. 2004. “Georeferencing of Museum Collections: A Review of Problems and Automated Tools, and the Methodology Developed by the Mountain and Plains Spatio-Temporal Database-Informatics Initiative (Mapstedi),” 29.

Nagendra, Harini, and Elinor Ostrom. 2012. “Polycentric Governance of Multifunctional Forested Landscapes.” International Journal of the Commons 6 (2): 104–33. doi:10.18352/ijc.321.

National Science Foundation. 2015. Advancing Digitization of Biodiversity Collections Program Solicitation. NSF 15-576.

Nelson, Gil. 2014. “IDigBio: The US National Science Foundation’s National Resource forDigitization of Biological and Palobiological Collections.” In Geological Society of America, Annual Meeting, Vancouver, Canada.

Nelson, Gil, Patrick Sweeney, Lisa E. Wallace, Richard K. Rabeler, Dorothy Allard, Herrick Brown, J. Richard Carter, et al. 2015. “Digitization Workflows for Flat Sheets and Packets of Plants, Algae, and Fungi.” Applications in Plant Sciences 3 (9): 1500065. doi:10.3732/apps.1500065.

Office of the Inspector General. 2017. “Final Evaluation Report – Evaluation of USGS Scientific Collection Management Policy, Report No. 2016-ER-057.” U.S. Department of the Interior, September, 7.

Ostrom, Elinor. 1990. Governing the Commons: The Evolution of Institutions for Collective Action. Cambridge University Press.

Ostrom, Elinor. 2008. “Polycentric Systems as One Approach for Solving Collective-Action Problems.”

Ostrom, Elinor. 2010. “Beyond Markets and States: Polycentric Governance of Complex Economic Systems.” American Economic Review 100 (3): 641–72. doi:10.1257/aer.100.3.641.

Ostrom, Elinor, Roy Gardner, James Walker, James M. Walker, and Jimmy Walker. 1994. Rules, Games, and Common-Pool Resources. University of Michigan Press.

Page, Lawrence M., Bruce J. MacFadden, Jose A. Fortes, Pamela S. Soltis, and Greg Riccardi. 2015. “Digitization of Biodiversity Collections Reveals Biggest Data on Biodiversity.” BioScience 65 (9). Oxford Academic: 841–42. doi:10.1093/biosci/biv104.

Page, Roderic D. M. 2013. “BioNames: Linking Taxonomy, Texts, and Trees.” Peerj 1 (October): e190. doi:10.7717/peerj.190.

Paul, Deborah, Austin R. Mast, Greg Riccardi, and Gil Nelson. 2013. “IDigBio as a Resource for the Digitization of a Billion Biodiversity Research Specimens.” In TDWG 2013 ANNUALCONFERENCE.

Poteete, Amy R., Marco A. Janssen, and Elinor Ostrom. 2010. Working Together: Collective Action, the Commons, and Multiple Methods in Practice. Princeton University Press.

Ruch, Jeff. 2018. “Information Correction Request Submitted under USGS Information Quality Guidelines.” Public Employees for Environmental Responsibility.

Ruch, Jeff. 2019. “Appeal to the USGS Response to a Request for Correction of Information Submitted under USGS Information Quality Guidelines.” Public Employees for Environmental Responsibility.

Samuelson, Paul A. 1954. “The Pure Theory of Public Expenditure.” The Review of Economics and Statistics 36 (4). The MIT Press: 387–89. doi:10.2307/1925895.

Schneider, Julio V., Renate Rabenstein, Jens Wesenberg, Karsten Wesche, Georg Zizka, and Jörg Habersetzer. 2018. “Improved Non-Destructive 2D and 3D X-Ray Imaging of Leaf Venation.” Plant Methods 14 (January). doi:10.1186/s13007-018-0274-y.

Scott, Carol. 2006. “Museums: Impact and Value.” Cultural Trends 15 (1). Routledge: 45–75. doi:10.1080/09548960600615947.

Selwood, Sara. 1999. “Access, Efficiency and Excellence: Measuring Non-economic Performance in the English Subsidised Cultural Sector.” Cultural Trends 9 (35). Routledge: 87–137. doi:10.1080/09548969909365090.

Skarbek, David. 2011. “Governance and Prison Gangs.” American Political Science Review 105 (4): 702–16. doi:10.1017/S0003055411000335.

Skarbek, David. 2016. “Covenants without the Sword? Comparing Prison Self-Governance Globally.” American Political Science Review 110 (4). Cambridge University Press: 845–62. doi:10.1017/S0003055416000563.

Snow, Neil. 2005. “Successfully Curating Smaller Herbaria and Natural History Collections in Academic Settings.” BioScience 55 (9): 771–779.

Snyman, Sandy J., Dennis M. Komape, Hlobisile Khanyi, Johnnie van den Berg, Dirk Cilliers, Dyfed Lloyd Evans, Sandra Barnard, and Stefan J. Siebert. 2018. “Assessing the Likelihood of Gene Flow From Sugarcane (Saccharum Hybrids) to Wild Relatives in South Africa.” Frontiers in Bioengineering and Biotechnology 6. Frontiers. doi:10.3389/fbioe.2018.00072.

Stanziola, Javier. 2008. “Developing a Model to Articulate the Impact of Museums and Galleries: Another Dead Duck in Cultural Policy Research?” Cultural Trends 17 (4). Routledge: 317–21. doi:10.1080/09548960802615455.

Stringham, Edward Peter. 2015. Private Governance: Creating Order in Economic and Social Life. 1st edition. Oxford; New York: Oxford University Press.

Suarez, Andrew V., and Neil D. Tsutsui. 2004. “The Value of Museum Collections for Research and Society.” BioScience 54 (1): 66–74. doi:10.1641/0006-3568(2004)054[0066:TVOMCF]2.0.CO;2.

Tarko, Vlad. 2015. “Polycentric Structure and Informal Norms: Competition and Coordination within the Scientific Community.” Innovation: The European Journal of Social Science Research 28 (1): 63–80. doi:10.1080/13511610.2014.985194.

Taylor, John W., and Eric C. Swann. 1994. “DNA from Herbarium Specimens.” In Ancient DNA, 166–181. Springer.

Thiel, Andreas. 2017. “The Scope of Polycentric Governance Analysis and Resulting Challenges.” Journal of Self-Governance and Management Economics 5 (3). Addleton Academic Publishers: 52–82.

Thiers, B. 2019. Index Herbariorum: A Global Directory of Public Herbaria and Associated Staff. New York Botanical Garden’s Virtual Herbarium.

United States Geological Survey. 2019. USGS Scientific Working Collections Management. IM CSS 2019-01. https://www.usgs.gov/about/organization/science-support/survey-manual/im-css-2019-01.

Wang, Wei, Shuo Yu, Teshome Megersa Bekele, Xiangjie Kong, and Feng Xia. 2017. “Scientific Collaboration Patterns Vary with Scholars’ Academic Ages.” Scientometrics 112 (1): 329–43. doi:10.1007/s11192-017-2388-9.

